# Whole genome sequencing and comparative genomic analysis reveal novel allelic variations unique to a purple colored rice landrace (*Oryza sativa ssp. indica* cv. Purpleputtu)

**DOI:** 10.1101/536326

**Authors:** V. B. Reddy Lachagari, Ravi Gupta, Sivarama Prasad Lekkala, Lakshmi Mahadevan, Boney Kuriakose, Navajeet Chakravartty, A. V. S. Krishna Mohan Katta, Sam Santhosh, Arjula R. Reddy, George Thomas

## Abstract

Purpleputtu (*Oryza sativa ssp. indica cv.* Purpleputtu) is a unique rice landrace from southern India that exhibits predominantly purple color. This study reports the underlying genetic complexity of the trait and associated domestication and de-domestication processes during its coevolution with present day cultivars. Along-with genome level allelic variations in the entire gene repertoire associated with purple, red coloration of grain and other plant parts. Comparative genomic analysis of the whole genome sequence of Purpleputtu (PP) revels total of 3,200,951 variants including 67,774 unique variations were observed in PP when compared with 108 rice genomes. Multiple sequence alignment uncovered a 14bp deletion in *Rc* (*Red colored,* a transcription factor of bHLH class) locus of PP, a key regulatory gene of anthocyanin biosynthetic pathway. Interestingly, this deletion in *Rc* gene is a characteristic feature of the present-day white pericarped rice cultivars. Phylogenetic analysis of *Rc* locus revealed a distinct clade showing proximity to the progenitor species *rufipogon* and *nivara.* In addition, PP genome exhibits a well conserved a 4.5Mbp region on chromosome 5 that harbors several loci associated with domestication of rice. Further, PP showed 1,387 unique SNPs compared to 3,024 lines of rice (SNP-Seek database). The results indicate that PP genome is rich in allelic diversity and can serve as an excellent resource for rice breeding for a variety of agronomically important traits such as disease resistance, enhanced nutritional values, stress tolerance and protection from harmful UV-B rays.

## Introduction

Rice is the staple food for more than half of the world’s population and substantially meets both food and calorie requirements. Rice cultivation covers about 165 million hectares globally with an annual production of 758.8 million MT (FAO, 2017) and is a critical component of the global food security system. Diverse Asian populations rely on rice to cover 35-80% of their calorie needs, while global reliance is about 21%. The two subspecies of cultivated rice, *O. sativa*, namely *indica* and *japonica*, occupy more than 90% of the Asian rice crop acreage (Muthayya *et al.*, 2014). Evolution of rice from its progenitors is marked by great complexity as it appears to have involved diverse lineages, domestication/de-domestication processes and selection, both natural and artificial. Domestication and selection of different populations of Asian wild progenitor rices, *O. rufipogan* and *O. nivara* might have begun more than 10,000 years ago giving rise to the present day Asian cultivated rices (Choi *et al.*, 2017; Qiu *et al.*, 2017; Yang *et al.*, 2015). Wild rices predominantly exhibit varying grain colours and this trait is known to be associated with domestication (Civáň and Brown, 2017). Rice germplasm collections comprise various colored rice lines, though these are neither cultivated widely nor used extensively in crop improvement programs. Colored rices have been widely used as entries in trials for the discovery of genes that confer resistance to bacteria, fungi and insects (Ahuja *et al.*, 2010). Coloured rices of various hues were described as red, brown, purple and black, based largely on pericarp and or hull colouration due to accumulation of anthocyanins, their precursors, flavonoids or their combinations, called co-pigmentation, besides other polyphenolic derivatives. Anthocyanins, the end products of anthocyanin pathway, are ubiquitous pigments known to be present in flowering plants. Naturally occurring rice landraces that accumulate anthocyanins, proanthocyanins and anthocyanin derivatives have been widely described (Reddy *et al.*, 1995; Oh *et al.*, 2018). Historically, colored rices have been deemed specialty rices by various ancient Asian cultures. For example, black rice has been described as forbidden rice or Emperor’s rice in China and red rices have been used in some religious celebrations in south and southeast Asia. However, due to changed consumer preference for white grained rices, they were not exploited in the breeding programs despite their special features such as enhanced levels of antioxidant compounds and biotic and abiotic stress tolerance (Reddy *et al.*, 2007). In addition, red/purple rices exhibit some well described domestication related traits, though in varying intensity, such as seed dormancy, grain shattering, photo-period sensitivity, long duration, tillering and lodging.

Purpleputtu (PP) is a colored landrace that exhibits purple colour in all aerial parts including seeds except in nodes and pollen (Reddy *et al.*, 1995). It is an *indica* landrace cultivated in small restricted areas in farmer fields in southern India, often used as border lines to demarcate test plots in experimental fields, primarily serving as a pollen barrier due to its height (Rangaswamy *et al.*, 1988). The genetic control of pericarp colour in PP has been described and molecular biological basis of the control of the underlying anthocyanin pathway has been elucidated (Reddy *et al.*, 1994; Reddy *et al.*, 1995; Reddy *et al.*, 2007; Oh *et al.*, 2018). Earlier studies on color in *japonica* rices revealed the contours of the genetic circuitry that govern color pathway (Furukawa *et al.*, 2007). Regulation of the anthocyanin pathway, both in *indica* and *japonica* subspecies, by different classes of transcription factors and repressors have been identified and tissue specific expression of some of these genes deciphered. (Reddy *et al.*, 1995; Sweeney *et al.*, 2006; Rahman *et al.*, 2013).

Allelic variations at certain target loci of the anthocyanin pathway that lead to the formation of many diverse flavonoids and anthocyanins have been described (Reddy *et al.*, 1995; Kim *et al.*, 2011; Maeda *et al.*, 2014; Kim *et al.*, 2015; Chin *et al.*, 2016). However, not much is known about allelic variations at loci associated with the pathway in terms of mutations, deletions and rearrangements. Scant information exists on differences at the genomic level between colored and white grained rices. Advancement of next generation sequencing (NGS) technologies along with the availability of the reference genome sequences for both *japonica* and *indica* rices provided an unprecedented opportunity to investigate the genome wide distribution of allelic variations that control complex pathways such as those that differentiate coloured rices from white ones. Deep sequencing coupled with comparative genomic analysis using extensively sequenced diverse rice lines and global rice data bases provide a great opportunity to gain incisive insights into the genetic and molecular basis of a diverse array of traits. Further, genomic analysis of diverse genotypes such as wild progenitors, land races, cultivars and modern rices is expected to throw new light on domestication, selection sweeps of specific genomic regions and evolution of colorless grain phenotype. Present day colored rices i.e. PP, had evolved from their colored progenitors such as *rufipogon* over thousands of years of cultivation, domestication and natural selection. It is interesting to investigate as to how such a complex trait governed by many genes across the genome has evolved and maintained even when selection is biased towards white grained rices in modern times. Interestingly, colored rices are known for traits associated with disease resistance and abiotic stress tolerance as demonstrated by numerous reports of introgression breeding via wide hybridisation to essentially transfer useful genes from progenitors and wild relatives to present day cultivars, a form of de-domestication.

The present study is aimed at understanding the basis of existence and maintenance of PP, a fully colored rice, by whole genome deep sequencing and comparative genomics using a global collection of thousands of rice lines including progenitors such as *rufipogan, nivara*; and the reference genome of *Nipponbare* rice (Kawahara *et al.*, 2013). We uncovered a significant number of genome-wide allelic variations in PP including those in genes associated with anthocyanin biosynthesis and genomic regions associated with the domestication-related genes controlling dormancy, seed shattering and diseases response. Additionally, we report here the discovery of novel alleles in genomic regions showing extreme conservation through evolution and thus representing selection sweeps. Besides, we identified numerous unique alleles at loci associated with major structural and regulatory genes of the pathways determining purple phenotype.

## Results

### Whole genome sequencing of PP and comparison with Nipponbare reference genome

Whole genome shotgun sequencing of PP genomic DNA (80x coverage) on Illumina HiSeq2000 platform yielded 43.47 GB of raw data that include a total of 430,403,016 paired end reads of 100bp. More than 81.7% of the data exceeded Q30 Phred quality score. The quality score of read 1 (forward) and read 2 (reverse) are shown in Supplementary Figure S1A & S1B respectively, while position-based quality of each nucleotide for read 1 & 2 is shown in Supplementary Figure S1C & S1D respectively. Read based GC content estimates show that more than 36% of reads have less than 30% of GC content (Supplementary Figure S1E). The data was deposited in NCBI SRA database with an accession number PRJNA309223. The reads were aligned to *japonica* rice cultivar MSU7 reference genome using BWA. Overall, 95.7% of the total reads were mapped covering 94% of the reference genome. The reads with mapping quality value ≥30 and insert-size between 100bp and 1Kb were retained for further analysis after removing duplicates. Out of total generated reads, 263,679,866 were aligned with reference genome and having average of 49.5X read depth and 80.61% genome-wide coverage (Table 1 and Supplementary Figure S2A). Supplementary Figures S2B, S2C and S2D indicate percentage of aligned reads, chromosome-wise average read depth and coverage respectively.

**Table 1:**
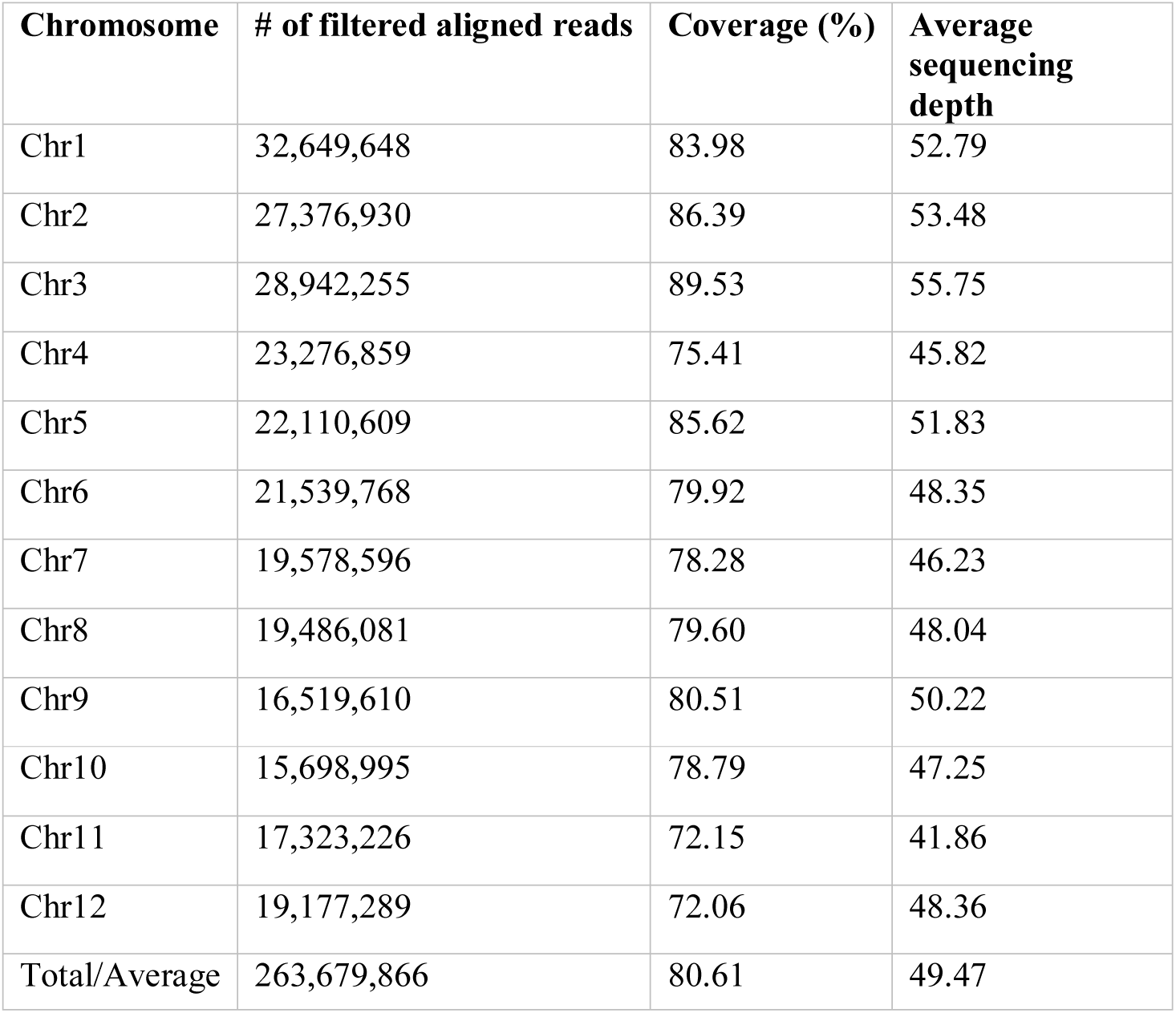
Chromosome wise distribution and read statistics of purpleputtu (PP) genome.

### Annotation of unmapped reads of PP genome

Assembled unmapped genome of PP was used for gene masking using rice as reference model. A total of 2,124 genes were predicted, out of which 1,239 gene sequences were annotated based on UniProt, NCBI NR and Phytozome database (Supplementary Table S1). Of the 1,239 genes sequences, set of 70 and 1069 have characterized and uncharacterized gene information annotated from homologous rice cultivar sequences. The remaining 76 sequences were annotated to the orthologous sequences of maize, wheat, purple false brome, cutgrasses and sorghum. A set of 885 sequences do not find suitable match in any of the above-said databases, indicating that they are unique in PP genome and can be denoted as uncharacterized sequences from PP.

### Identification of variants in PP genome and variant desert regions

Comprehensive genome-wide mapping diagram indicates the read depth, gene density, insertion density, deletion density and SNP density (Figure 1). A total of 3,200,951 variants (2,824,513 SNPs and 376,445 INDELs) were identified in PP (Table 2) with read depth ≥ 5 and variant quality score ≥ 50 against Nipponbare reference genome (Supplementary Table S2). A majority of the variants (88.24% of SNPs and 94.44% of INDELs) were found to be homozygous. Most of the SNP changes observed were of transition type: A > G (18.92%), C > T (16.47%), G > A (16.43%) and T > C (18.89%) with a Ts/Tv ratio of 2.41 (Figure 2A). A majority (70%) of the identified changes are short INDELs of length 1-2bp (Figure 2B). Of the total variants 1,058,815 (33.8%) map to the repeat region of the genome (Supplementary Table S3). The SNP density was estimated to be 756 SNPs and 100 INDELs per 100 Kb in PP in comparison with MSU7 assembly. Chromosome (Chr) 10 shows the highest SNP density (889/100Kb), while the lowest SNP density (624/100 Kb) was found in Chr5. Chr1 shows the highest INDEL density (109/100 Kb) whereas Chr4 has the lowest (85/100Kb) (Supplementary Table S2). We observed that 67% of the variant desert region (Chr5) falls into repeat regions with a majority of the repeat class being putative retrotransposons as compared with a few other *indica* lines (Wang *et al.*, 2009). The average read depth and coverage in this variant desert region was observed to be 43X and 81%. A total of 652 genes overlap were found in the variant desert region, half of which do not have a single variant. Of these genes 368 and 43 genes belong to the retrotransposon and transposon proteins respectively (Supplementary Table S4 & S5).

**Figure 1:**
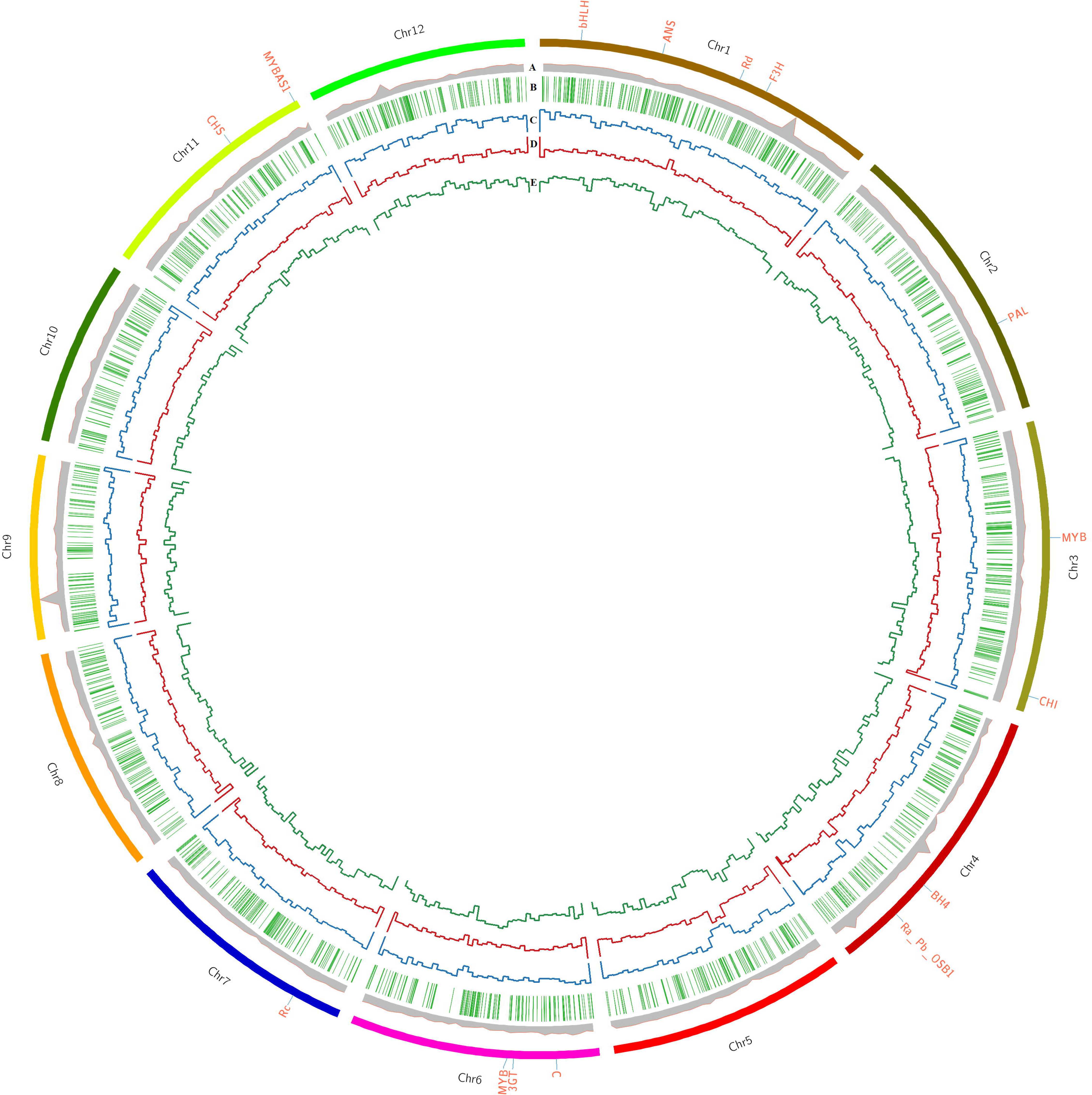
Comprehensive genome-wide analysis of Purpleputtu (PP) characteristics with Nipponbare reference genome. Circle diagram represents (A) Read depth (B) Gene density (C) Insertion density (D) Deletion density (E) SNP density. Outer circle of diagram represents the chromosomes of PP genome, each one labeled with important genes.

**Figure 2:**
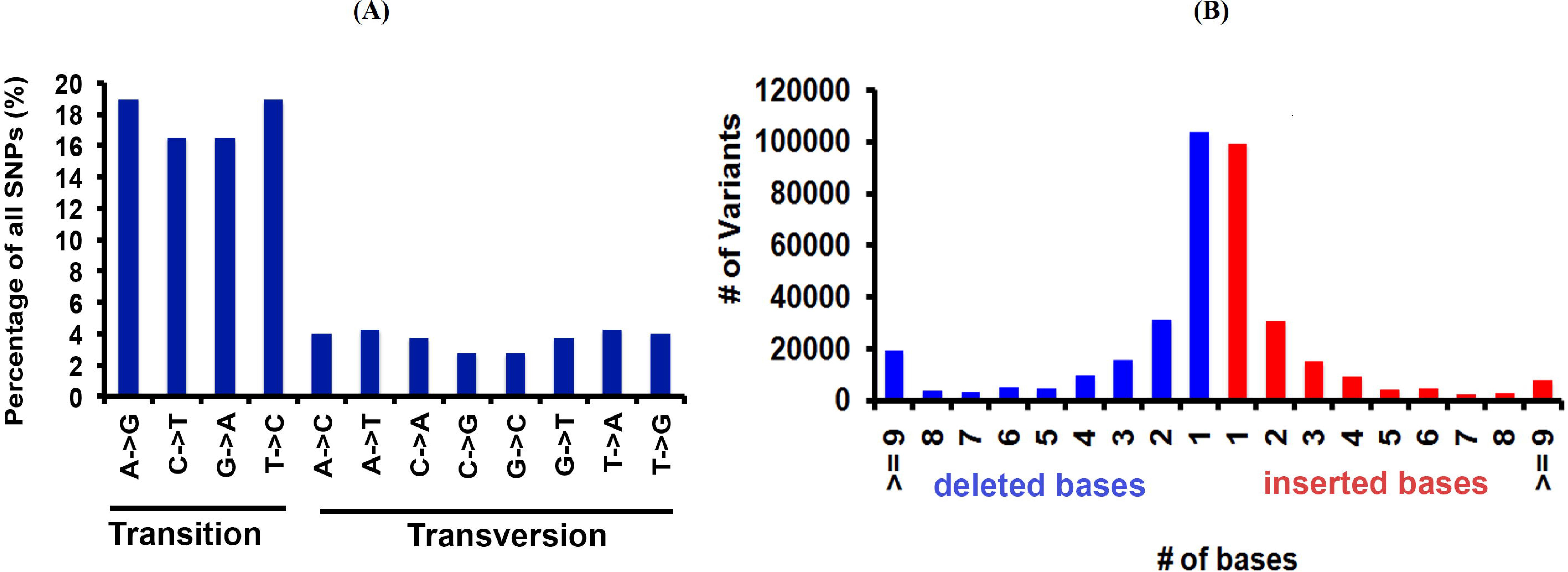
Base distribution statistics of variants identified in PP genome. (A) Percentage of transition and transversions (B) Distribution of deleted and inserted bases.

**Table 2:**
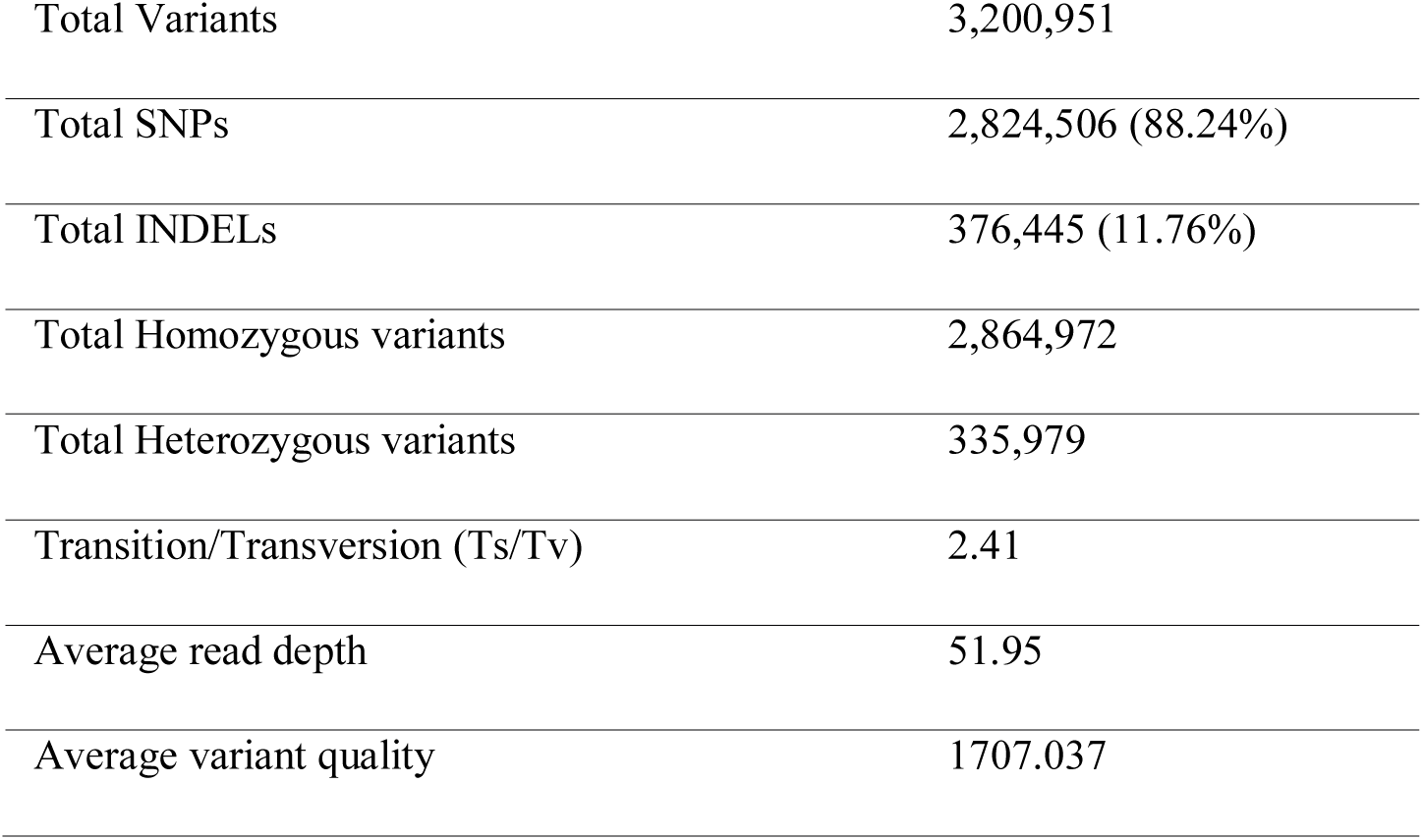
Summary statistics of various identified variants.

### Annotation of variants in PP genome

The variants were annotated using an in-house pipeline against the gene model provided by MSU7 database. Overall, ∼32% of the variants span the genic region and the remaining 68% fall in the non-genic regions (Figure 3A). Out of the total genic variants, 50% overlap exonic region and of these 79.6% falls in the coding regions. Of the total variants in the coding regions, 37.6% were synonymous and the remaining 63.4% were non-synonymous type. About half (55.8%) of the non-synonymous type variants belong to missense class. We found that 33% of the total variants are in the repeat regions. Of these, 23.3% were genic and the remaining 76.7% were inter-genic. Further, a breakup of repeat class variants revealed that many of them belong to retrotransposons and few to Miniature inverted-repeat transposable element (MITE) repeat class and Cacta-like transposons (Figure 3B). These may serve as a valuable resource for selecting functional markers in genetic mapping programs. We further investigated the variant density around the transcription start sites (TSS) and found that it peaks at around 420bp upstream and dips at ∼165bp downstream (Figure 3C). The variant density gradually decreases towards zero on both sides of the TSS. Deep analysis of variants presents in 1.5Kb upstream shows that maximum variations were present in the genes for pyrrolidone-carboxylate peptidase, WD domain/G-beta repeat domain containing protein, actin, dehydrogenase E1 component domain containing protein and CBL-interacting protein kinase 1 (Figure 3D), all variants and associated genes are depicted in Supplementary Table S6.

**Figure 3:**
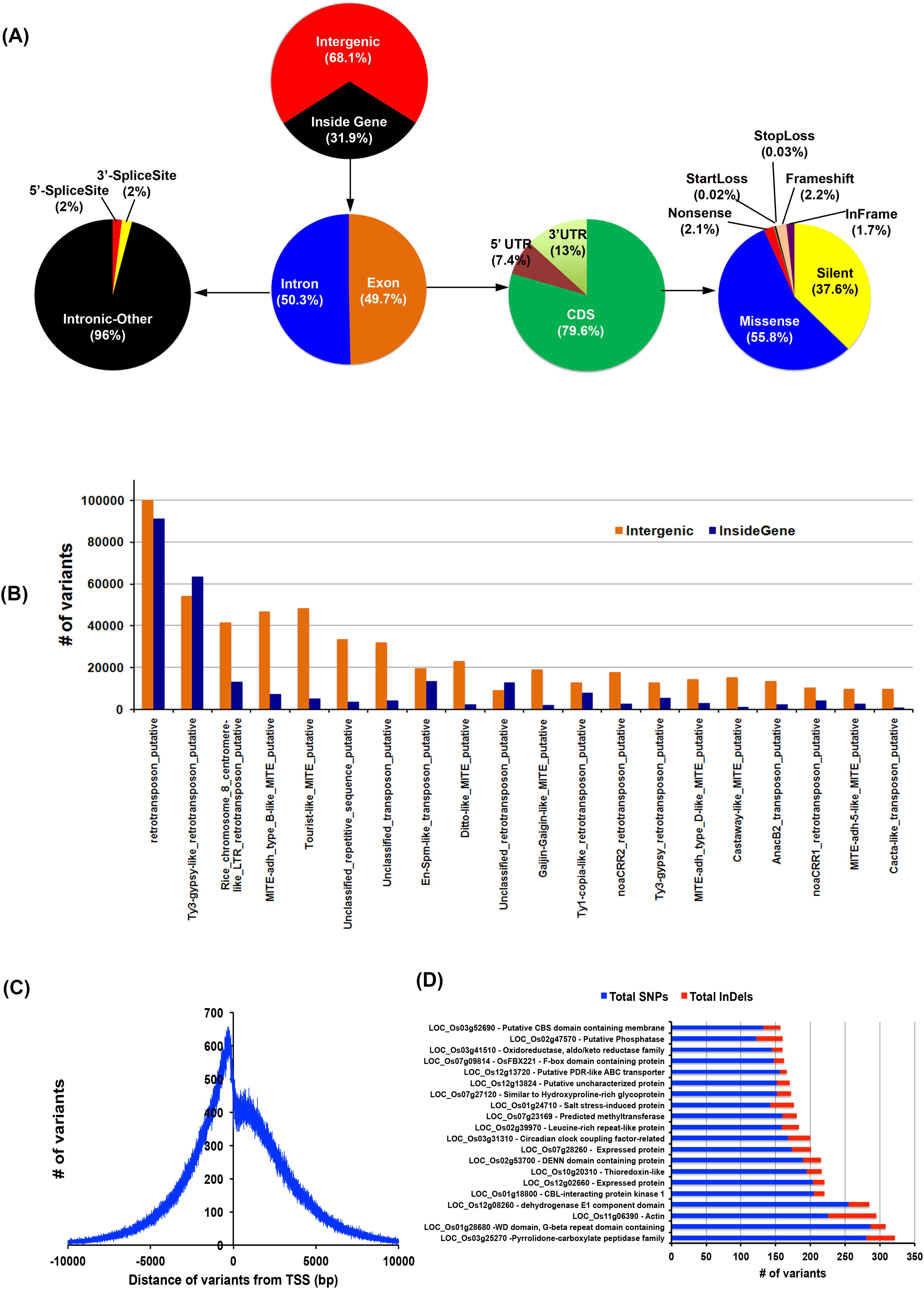
Comprehensive variant detection and annotation (A) Indicates the distribution of variants in different regions of genome (B) Represent the number of variant in different types of repeats (C) Variant density graph from transcription start site (TSS) and (D) Genes containing highest SNPs/INDELs variations.

### Unique variants of PP genome

To capture unique variants of PP, we compared its genome with a panel of 108 genome sequences (Supplementary Table S7) covering different red rices, progenitors, landraces, modern cultivars spanning across sativa group (*indica*, temperate and tropical *japonica*) and Australian, aromatic and wild rices. This panel also includes other wild/progenitor species such as *longistaminata, brachyantha, barthii, meridionalis, nivara, glaberrima* and *rufipogon*. A deeper analysis of the genome sequences with this panel allowed us to identify a set of novel and unique alleles in PP genome. Out of 3,200,951 variants, 67,774 were found to be unique to PP (Supplementary Table S8). Of these, 64,394 are SNPs and 3,380 are INDELs, which include 390 INDELs that span exonic regions. Zygosity analysis of unique variants revealed that 49,025 (72.74%) are homozygous and 18,749 (27.66%) are heterozygous. Among these unique variants, 24,576 were mapped to genic regions and 43,198 to intergenic regions (Supplementary Table S8). The genic regions (12,831) span across 9,370 genes indicating that unique variations occur in almost one fifth of the total genes in rice. Further classification of unique variants in 5’ UTRs, 3’ UTRs, intronic regions and splice junction sequences shows 7,087 missense SNPs, 4,942 silent variations, 371 nonsense mutations and 23 start-loss and 18 stop-loss variants. Analysis of SNP desert region of chromosome 5 revealed 224 unique variants of which 13 are INDELs and 211 SNPs (Supplementary Table S9). A set of 63 variants are uncovered in genic region of which 34 are silent mutations, 26 are missense mutations and one is a nonsense mutation. A majority of these variations are localized in Ty-3 gypsi class of retrotransposons. Variant analysis also identified 96 unique variants of PP genome associated with morphological traits, physiological traits and resistance or tolerance to biotic and abiotic stresses (Supplementary Table S10).

### Comparison of PP genome with global collection of 3,024 rice lines

PP genome was subjected to a deep comparative analysis by aligning with International Rice Genome Sequencing Project (IRGSP v1.0) assembly (http://rgp.dna.affrc.go.jp/IRGSP/) having 3 K SNP-Seek data set. The resultant 1,962,843 SNPs in PP were compared with 5,854,680 SNPs of 3 K global rice collection to identify rare variants of large effect. PP variants were merged with those of 3 K SNP-Seek dataset and rare variants were obtained with minor allele frequency (MAF) cutoff of <0.01. The dendrogram made with 3 K dataset comparison shows that PP is close to IRIS-313-8921, IRIS-313-8498 (Supplementary Figure S5). A total of 481,205 rare SNP loci were found amongst the combined dataset; which has 5,323,594 SNPs in common variant loci. Of all rare variants, 479,818 loci were only called in PP and not called in 3 K dataset; therefore, they were denoted as novel. The remaining 1,387 SNP loci/SNPs were found to be unique to PP (Supplementary Table S11). The chromosome-wise distribution of these unique SNPs are as follows: Chr1 (141), Chr2 (100), Chr3 (100), Chr4 (129), Chr5 (108), Chr6 (124), Chr7 (114), Chr8 (132), Chr9 (102), Chr10 (101), Chr11 (128), Chr12 (108). The localization of these SNPs in their respective loci is listed in Supplementary Table S11.

### Functional classification of variants

The functional classification of the variants was performed at different levels: pathway, ontology and traits. For the pathway study we compared the variants against rice metabolic pathway RiceCyc v3.3 database (Dharmawardhana *et al.*, 2013). The pathways with the highest number of variants include cytokinins glucoside biosynthesis, betanidin degradation, sucrose degradation to ethanol and lactate, and cellulose biosynthesis (Supplementary Table S12) (Gupta *et al.*, 2017a). Betanidin degradation eliminates Betacyanin pigment pathway (which leads to production of red, purple and violet betacyanin pigments which are predominant in *Caryophyllaceae*); here it is worth noting that anthocyanins and betacyanins are mutually exclusive in flowering plants (Rodriguez-Amaya, 2018). Similarly, sucrose degradation is a prerequisite for preventing root hypoxia. There is some information on the role of this process in salinity tolerance (Behr *et al.*, 2017). Cytokinin glucoside biosynthesis is reported to be associated with indeterminate growth. Of the total unique variants in PP, 25,447 variants mapped to genes associated with 338 pathways (Figure 4). Of these, 13,439 are silent, 10,871 are missense, 66 are nonsense, 26 are start-loss and 43 are stop-loss mutations. Interestingly, PP exhibits many unique variations at diverse loci controlling the highly conserved ubiquitous flavonoid biosynthetic pathway. These variants span across genes controlling sub-pathways such as flavonoid biosynthesis (PWY1F-FLAVSYN), flavonol biosynthesis (PWY-3101), anthocyanin biosynthesis [pelargonidin 3-O-glucoside, cyanidin 3-O-glucoside] (PWY-5125), anthocyanin biosynthesis [delphinidin 3-O-glucoside] (PWY-5153), proanthocyanidin biosynthesis from flavanols (PWY-641). Besides this, other biotic and abiotic stress responsive pathways were also found to harbour unique variants in PP (Supplementary Table S11) (Gupta *et al.*, 2017a; Gupta *et al.*, 2017b; Gupta *et al.*, 2018a). Specifically, 122 variations were identified in flavonoid biosynthesis pathway (PWY1F-FLAVSYN) genes of which 62 are silent and 57 are mis-sense variants, 1 each of nonsense, frame-shift and in-frame variations. As many as 40 variants were observed in the flavonol biosynthetic pathway (PWY-3101). Considering the pathway-based analysis, it is inferred that pelargonidin/cyanidin 3-O-glucoside sub-pathway (PWY-5125) consists of 34 variations in which 9 are silent, 23 are missense, 1 each of frame-shift and in frame variations. In contrast, 24 variations were observed in an evolutionarily silenced delphinidin route of anthocyanin biosynthesis pathway (PWY-5153) of rice. Only 1 missense mutation was observed in enzymes responsible for proanthocyanidin pathway from flavanols (PWY-641) in rice. Trait ontology analysis for unique variants in PP was performed using Q-TARO database (Yonemaru *et al.*, 2010). Out of the 67,774 variants, 6,283 were mapped to the regions associated with morphological (2,287), physiological (1,361), resistance or tolerance traits (2,436) and other traits (199) (Supplementary Table S13). Among these 2,812, 2, and 8 variants were found to be missense, start-loss and stop-loss respectively. Interestingly, the start-loss variations are observed only in flowering and cold tolerance traits whereas stop-loss variants were observed in genomic locations associated with sterility, and various biotic and abiotic stress traits such as blast resistance, soil stress tolerance, drought tolerance and cold tolerance (Gupta *et al.*, 2017b). Of all the variants that mapped to the trait-associated loci, the highest number of variants mapped to dwarf, sterility and blast resistance categories (Figure 5, Supplementary Table S13). In addition, variants were observed in traits related to seed, leaf, flowering, germination, drought tolerance, stress tolerance and lodging resistance which reflect the typical phenotype of PP.

**Figure 4:**
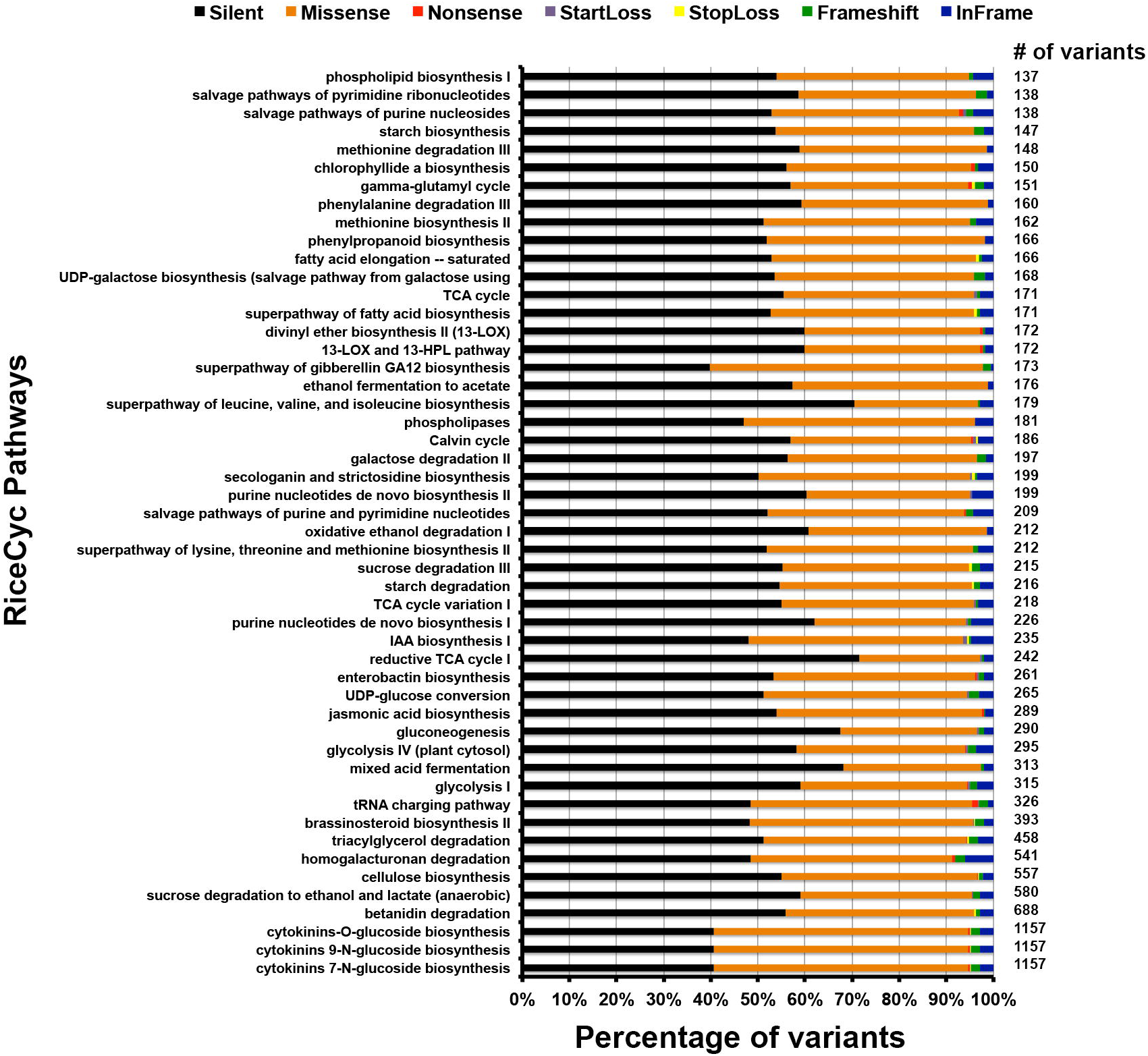
Bar diagram showing total number of different variants and their association in various metabolic pathways.

**Figure 5:**
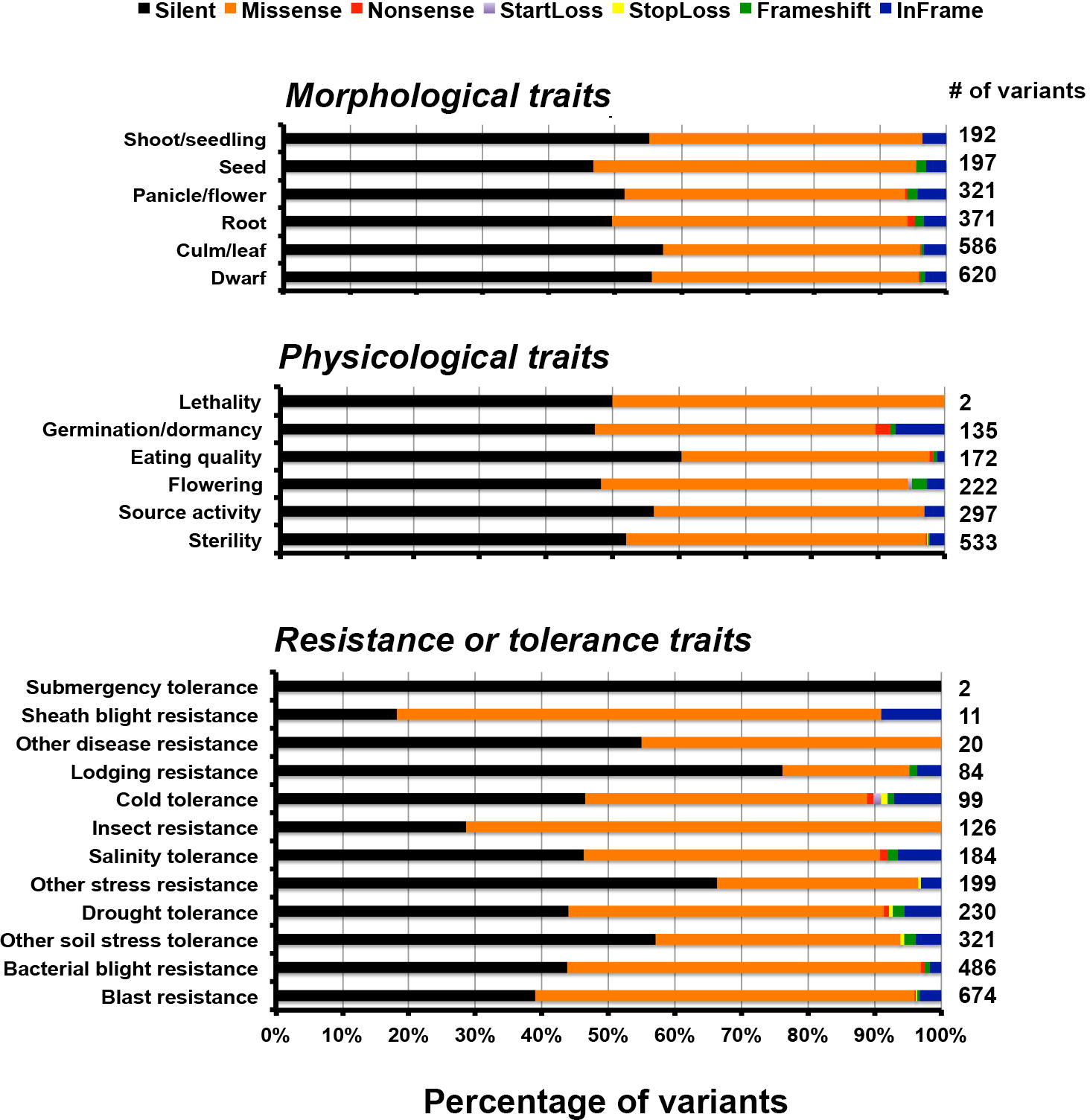
Bar diagram showing total number of different variants involved in morphological, physiological traits and resistance to biotic and abiotic stress.

### Variations in transcription factor genes

The unique variants of PP mapped to different transcription factors (TFs) in rice including MYB, bHLH, FAR1, NAC, WRKY, ERF, bZIP, and AP2 (Supplementary Table S14). The highest number of variants (1752) was localized in Far-Red Impaired Response1 (FAR1) family transcription factors which controls the far-red light signaling pathway by modulating phyA expression by activating FHY1 and FHL, thus maintaining homeostasis (Lin *et al.*, 2007; Gupta *et al.* 2019). In contrast, whirly family of transcription factors involved in defense response show low number of variants, viz., three, indicating high level of conservation. NAC transcription factor family, which is one of the largest families of plant-specific transcription factors and plays an important role in response to various plant stresses shows 682 variants (Olsen *et al.*, 2005; Nakashima *et al.*, 2012). The bHLH class of transcription factors, including the Rc (Red color), which are involved in flavonoid/anthocyanin biosynthesis besides other biological processes such as wound and drought stress, light signaling and hormone signaling shows 672 variants (Carretero-Paulet *et al.*, 2010). Interestingly, 319 of them are missense, 3 nonsense, 15 frame-shift and 79 in-frame mutations and the remaining are silent; all of these changes put together indicates significant variation in this locus. MYB genes and related genes of rice, known to be involved in the regulation of anthocyanin biosynthetic pathways as well as in other different biological processes in rice were found to have 898 variants of which 433 are missense variants (Liu *et al.*, 2015; Ambawat *et al.*, 2017). The WRKY class of transcriptional factors show 483 variants of which 157 are silent and 272 are missense mutations, which are involved in several biotic and abiotic stress responses (Phukan *et al.*, 2016, Gupta *et al.*, 2018b). A set of 47 missense variations and one stop-loss mutation is observed in AP2 transcription factor family genes which are involved in abiotic stress response in rice (Fu *et al.*, 2010).

### Evolutionary analysis of Bh4 domestication genes

We studied independent events of domestication in the PP genome, which can offer an extremely useful system for studying the genetic basis of parallel evolution with other rice genomes. A significant trait altered by rice domestication and de-domestication is hull color. Wild progenitors of two cultivated rice species have predominantly black-colored hulls and straw coloured hulls; these are the phenotypic effects of Bh4 and straw hull 4 (Sh4) candidate gene expression (Vigueira *et al.*, 2013; Zhu *et al.*, 2011). Examination of evolutionary relationship of Bh4 genes of PP with other 70 rice cultivar genes is shown in Supplementary Figure S3. The phylogenetic tree shows the Bh4 DNA sequence variations, that provides clues to the parallel evolution of hull color variation in PP and other rice cultivars. Supplementary Figure S3 shows that out of 70 rice cultivars, 19 have variations suggesting that the same gene is responsible for parallel trait evolution.

### Variations in Anthocyanin pathway genes

The variants that mapped to the anthocyanin pathway genes were analyzed for the resultant changes in the encoded proteins and these include both regulatory and structural genes such as C (Chr6), Ra (Chr4), Rc (Chr7), basic helix-loop-helix TF (bHLH) (Chr1), Rd (Chr1), MYB (Chr3), MYB (Chr6), MYBAS1 (Chr11), chalcone synthase (CHS) (Chr11), chalcone isomerase (CHI) (Chr3), leucoanthocyanidin dioxygenase (ANS) (Chr1), Flavanone 3-hydroxylase (F3H) (Chr1), 3-O-glucosyltransferase (3GT) (Chr6), black hull 4 (BH4) (Chr4) and Phenylalanine ammonia-lyase (PAL) (Chr2) (Figure 1). The variants of anthocyanin pathway genes were explored to understand the key differences that may explain the uniqueness of PP genome in showing distinct color phenotype (Cingolani *et al.*, 2012). Of the 585 variants observed, 388 were identified as upstream variants followed by intronic (99), missense (45), synonymous (37) and 1 stop-gain variant (Supplementary Table S15). Comparative analysis of 5cM Rc locus (Chr7) spanning 42.6 cM to 47.7 cM in 3 BAC clones (AP003748, AP005098, AP005779) revealed unique variations in PP placing it as a distinct clade within the *indica* group, separated from all other rice lines (Sweeney *et al.*, 2006). Contrary to expectations, PP with its colored pericarp phenotype shows the 14bp deletion signature in Rc gene which is a characteristic feature of the present white pericarped rice lines. Further evolutionary relationship of PP Rc gene to other Rc gene homologs of rice cultivars indicate divergence from other rice cultivars (Supplementary Figure S4). MYBAS1 gene (LOC_Os11g47460) on Chr11 was found to have 56 missense SNPs responsible for changes in 42 amino acids, which could be a possible paralogue of the Rc gene on Chr7. In addition to that, the Rc gene (LOC_Os07g11020) on Chr7 exhibits an intronic variation responsible for deletion of ‘GAGA’ at position 138 that does not affect the functions of the gene. In addition, homology search for Rc genes against other rice genomes, did not provide any significant hit for any alternate functional gene. Notably, anthocyanin regulatory gene Ra (Pb/OSB1/ LOC_Os04g47080) on Chr4 which is reported to be associated with purple color pericarp (Wang and Shu, 2007) shows one missense variation leading to M64T, and two frame shift variations causing T575fs and V545fs. Both of these frameshift mutations in this gene lead to alternate chain of 30 amino acids (PLGAGINIGWSPWTDTS QVCLICCRRTWE*) in the C terminus when compared to Nipponbare reference genome. bHLH (LOC_Os01g09900) on Chr1 shows interesting variations leading to amino acid changes (E304K, P148L, V76A and disruptive in-frame insertion at A98AA). MYB gene (LOC_Os06g19980) on Chr3 shows a total of 14 variations in which 7, 3, 2, and 1 are downstream gene variant, in-frame deletion, intron variant and synonymous variant, respectively; the remaining 1 variant belongs to missense variation and the changes in amino acid A263G possibly does not alter the secondary structure of the protein. MYB genes (LOC_Os06g19980) on Chr6 has 23 missense variations causing changes in amino acids *Viz.* S27R, M47L, R63L, R66L, D72G, L80N, I82S, A83P, I86 V, Q112E, S116I, E147G, E148D, I151 V, L163 V, T202A, I232 T, L296R, R298G, S446L, Y607H and A639P; these changes possibly alter the structure and function of this MYB protein. The gene encoding PAL (LOC_Os02g41630) enzyme on Chr2 catalyzing the formation of 4-coumaroyl-CoA from phenylalanine has 1 synonymous, 1 intronic variation and only one missense variation changing amino acid A621 V of the protein. CHS (Os11g0530600) on Chr11 has 1 SNP leading to a single amino acid change, N158S. F3H (LOC_Os01g50490) on Chr1, the gene encoding the enzyme involved in conversion of flavonone to dihydroflavonones has 49 unique variations in which 22 SNPs lead to protein structure variations. We found 4 SNPs each causing amino acid changes to proline and glutamic acid *viz*.T3P, A79P, L99P, L474P, and Q80E, K407E, D410E, D414E respectively. In addition, the remaining SNPs cause mutations of L6V, T15M, V34A, S55G, N77T, D103N, A110 V, C389S, L392F, I393V, R418G, G420D, G424W, and T488A.

Possibly, these mutations contribute towards hyper-accumulation of anthocyanins in PP (Han *et al.*, 2010). 1 silent variation was identified in ANS1 (LOC_Os01g27490) gene on Chr1, encoding the key enzyme involved in the conversion of leucoanthocyanidins to anthocyanidins, the penultimate step in anthocyanin biosynthesis. The last enzyme 3GT (LOC_ Os06g18790) of this pathway was found on Chr6 of PP genome, converting anthocyanidins to anthocyanins and had 3 unique SNPs (M186 V, K190Q, K190R) that may have a role in accumulation of pigments in PP (Brazier-Hickes *et al.*, 2007).

## Discussion

*Oryza sativa*, an independently domesticated rice that has been in cultivation for more than 12,000 years, has become the predominant cereal staple for most of the ∼3 billion plus Asians. With an estimated 30% rise in population by 2050 and unpredictable climate changes, rice breeders must gear up for substantial increase in rice productivity across various agroclimatic regions. Modern rice breeding technologies are increasingly utilizing the genetic resources of progenitors and wild relatives by introgression of new genes associated with important agronomical traits into cultivars. These mainly include genes for biotic and abiotic stress tolerance, growth and maturity traits. Though colored rices constitute a significant proportion of germplasm collections, they were not extensively used as genetic resources in breeding. Progenitors and domesticated colored rices were reported to be potential source of genes for resistance to diseases and pest, and tolerance to abiotic stresses. Besides, diverse colored pigments are recognized for their nutritional quality and anti-oxidant properties. Further, colored rices were found to be good subject material for understanding domestication and de-domestication processes in grasses in general and rice in particular (Choi *et al.*, 2017; Qiu *et al.*, 2017). We set out to uncover novel and unique alleles in one such fully colored rice line, PP, that is cultivated sporadically with no evidence of any directional selection or crossing in its long history of domestication and cultivation.

Whole genome sequencing of PP rice and mapping with Nipponbare genome was performed to discover genome wide DNA variations. A total of 263,679,866 reads were mapped to the reference genome. The assembly and annotation of unmapped reads indicates the presence of many uncharacterized genes having homology with the other wild species of *Oryza* such as *nivara, meridionalis, glumipatula, glaberrima, brachyantha, barthii, punctata, rufipogan* and *alta* (Supplementary Table S1). Interestingly, 56 gene models showed homology with a red rice line (*Oryza punctata)* and all these genes are uncharacterized indicating the presence of novel genes associated with red/purple pigmentation. Further, comparative genomic analysis using a wide variety of rice lines (108 panel) spanning both *indica* and *japonica*, progenitors, land races uncovered 64,349 SNPs and 3,380 INDELs unique to PP genome. In all, we captured unique variants in one fifth of the total gene models of rice. Of the 3,200,951 polymorphic SNPs identified, about a third span across exonic regions and three fourth of them fall in coding sequences. Further, about 33% of the variants are mapped to repeat regions, retrotransposons and transposable elements, a finding that falls within the range reported for *Oryza* (Stein *et al.*, 2018). The present data show that Chr1 has the highest number of INDELs and Chr10 has the highest number of SNPs per 100 kb. Similarly, the lowest INDEL density was on Chr4 and lowest SNP density on Chr5. In addition, distribution of SNP-rich and SNP-poor regions in each chromosome of PP was identified, which also corroborates with earlier findings in rice, Arabidopsis, and wheat (Subbaiyan *et al.*, 2012; Nordborg *et al.*, 2005; Ravel *et al.*, 2006).

The mapped genome of PP revealed a clear bias towards transitions (almost twice that of transversions) deviating from the expected ratio of 0.5. Higher Ts/Tv ratios were reported in rice, maize, otus, medicago, diploid wheat, *Triticum monococcum* and barley (Bindusree *et al.*, 2017; Subbaiyan *et al.*, 2012; Batley *et al.*, 2003; Vitte *et al.*, 2006). Due to wobble effect, transitions manifest mostly into silent mutations that do not alter the amino acid and thus conserves the amino acid chain (Wakeley, 1996). Among transitions, the C/T transitions were more in number, presumably due to a simple methylation being the cause of this mutation (Coulondre *et al.*, 1978). Higher frequency of C/T mutations has been reported in other crops such as common bean, maize, grape and citrus (Ramírez *et al.*, 2005; Batley *et al.*, 2003; Lijavetzky *et al.*, 2007; Terol *et al.*, 2008). Generally, genomic segments with higher SNP frequency were shorter than the lower SNP frequency segments. An SNP-poor region having 4.3Mb between 9.3Mb and 13.6Mb on Chr5 was identified in PP genome that is popularly described as ‘SNP desert’ and described in detail (Supplementary Table S4, S5 & S9). This conserved region in rice has been earlier reported in certain *indica* and *japonica* lines (Subbaiyan *et al.*, 2012; Wang *et al.*, 2009; Nagasaki and Yamamoto, 2010). It is well known that selective sweeps during the long process of domestication in both *indica* and *japonica* rice are responsible for lower SNP frequencies across some regions in the genome (Subbaiyan *et al.*, 2012; Caicedo *et al.*, 2007).

The progenitors of the present-day rice lines and most of the wild rices exhibit red color in the pericarp and some other plant parts. However, the present-day modern rice cultivars predominantly have white pericarp. The disappearance of purple-red-black-brown color in grain in modern cultivars is a great example of agronomical spreading of a single variation at the Rc (Red color) locus through the course of domestication and natural and artificial selection (Furukawa *et al.*, 2007; Sweeney *et al.*, 2007; Gross *et al.*, 2010). The Rc locus of PP shows the well characterized 14bp deletion in the fifth exon; this deletion has been reported to be the hallmark of white rices, yet the pericarp of PP that bears this signature, is colored. In contrast, almost all white pericarped lines, including all modern rice cultivars, have the same deletion at Rc locus. The Rc gene encodes a bHLH TF that regulates DFR activity. We predict an alternate TF that may be regulating DFR in PP grain. It is to be noted that the Rc locus in the colored progenitor *rufipogon* lacks this 14bp deletion. It may be the result of gene flow from cultivars to PP which indicates de-domestication. Introduction of many disease resistance genes, such as Xa21, from wild progenitors by introgression breeding and directed selection is a routine approach to exploit novel genes. We conclude that the Rc locus of PP is conserved due to simple cultivation and non-directional selection for colour. The white pericarp phenotype controlled by the deletion-containing Rc locus in modern rice cultivars is due to selection pressure for white grain. Our results challenge the notion that the 14bp deletion is an invariant signature of white rices only.

In depth analysis of variants revealed several unique SNPs/INDELs in structural and regulatory genes of various pathways responsible for stress response, genotypic and phenotypic effects on different traits of PP rice (Supplementary Table S12 & S13, Figure 4). Loci associated with cytokinin glucoside biosynthesis, betanidin degradation, sucrose degradation to ethanol and lactate, and cellulose showed the highest number of variants. Furthermore, our results revealed unique variations at loci associated with biosynthesis of betacyanin, anthocyanins and flavonoids. It is well known that betanidin degradation is a prerequisite step in eliminating betacyanin production in plants. It is worth noting that most flowering plants do not accumulate betacyanins and thus these two different color pigments are mutually exclusive (Rodriguez-Amaya, 2018; Shoeva *et al.*, 2016, Khoo *et al.*, 2017). Sucrose degradation is a prerequisite for preventing root hypoxia and salinity tolerance in some plants. Cytokinin glucoside biosynthesis is reported to be associated with indeterminate growth. The unique variant analysis uncovered 25,447 variants mapped to different genes associated with 338 pathways (Figure 4). Many structural, regulatory and inhibitory genes (Table 3) dispersed across the genome are involved in the anthocyanin pathway that leads to purple color formation in various tissues in PP (Reddy *et al.*, 1995). The production of anthocyanins such as cyanidin, pelargonidin, and delphinidin derivatives through a multistep anthocyanins pathway is mediated by several enzymes Phenylalanine ammonia lyase (PAL), cinnamate 4-hydroxylase (C4H), 4-coumarate CoA ligase (4CL), CHS, CHI, F3H, dihydroxyflavonol reductase (DFR); leucoanthocyanidin dioxygenase (LDOX)/anthocyanidin synthase (ANS), GT/3GT, acyltransferase (AT), methyltransferase (MT) (Ban-Simhon *et al.*, 2011). We found several unique variations in genes encoding PAL, CHS, F3H, ANS and 3GT enzymes of this pathway in PP when compared with 108 and 3 K reference databases. Variation (A621V) was found in alpha helical region of the PAL enzyme that is positioned far away from the catalytic site, indicating unlikely effect on structural and functional role of this enzyme (Bata *et al.*,2018). Changes of N158S in CHS enzyme in PP when compared with CHS crystal structure of *Freesia hybrida* (PDB ID:4WUM) indicated change of charged side change amino acid N to uncharged side chain amino acid to S at a location near the catalytic residues in PP CHS. This change may be enhancing the catalytic activity of the enzyme (Sun *et al.*,2015). The hydroxylation pattern of flavonoids controls their color, stability, and antioxidant capacity. We identified the gene encoding F3H in PP having 22 unique mutations, when compared with 3 K dataset of rice, which may be relevant in the maintenance of purple color through generations. These may also be relevant for stability, antioxidant activity and stress defense capacity (Liu *et al.*, 2014). Furthermore, analysis of structural and functional differences of PP F3H enzyme with that of white rice and other related cultivars indicates that it belongs to the plant Cytochrome P450 gene family (Schoenbohm *et al.*, 2000). Similarly, several silent mutations were identified in the *Ans* gene encoding ANS1 enzyme that catalyzes the penultimate step of the anthocyanin pathway, namely conversion of leucoanthocyanidin to anthocyanidin (Reddy and Reddy, 2010).

**Table 3:**
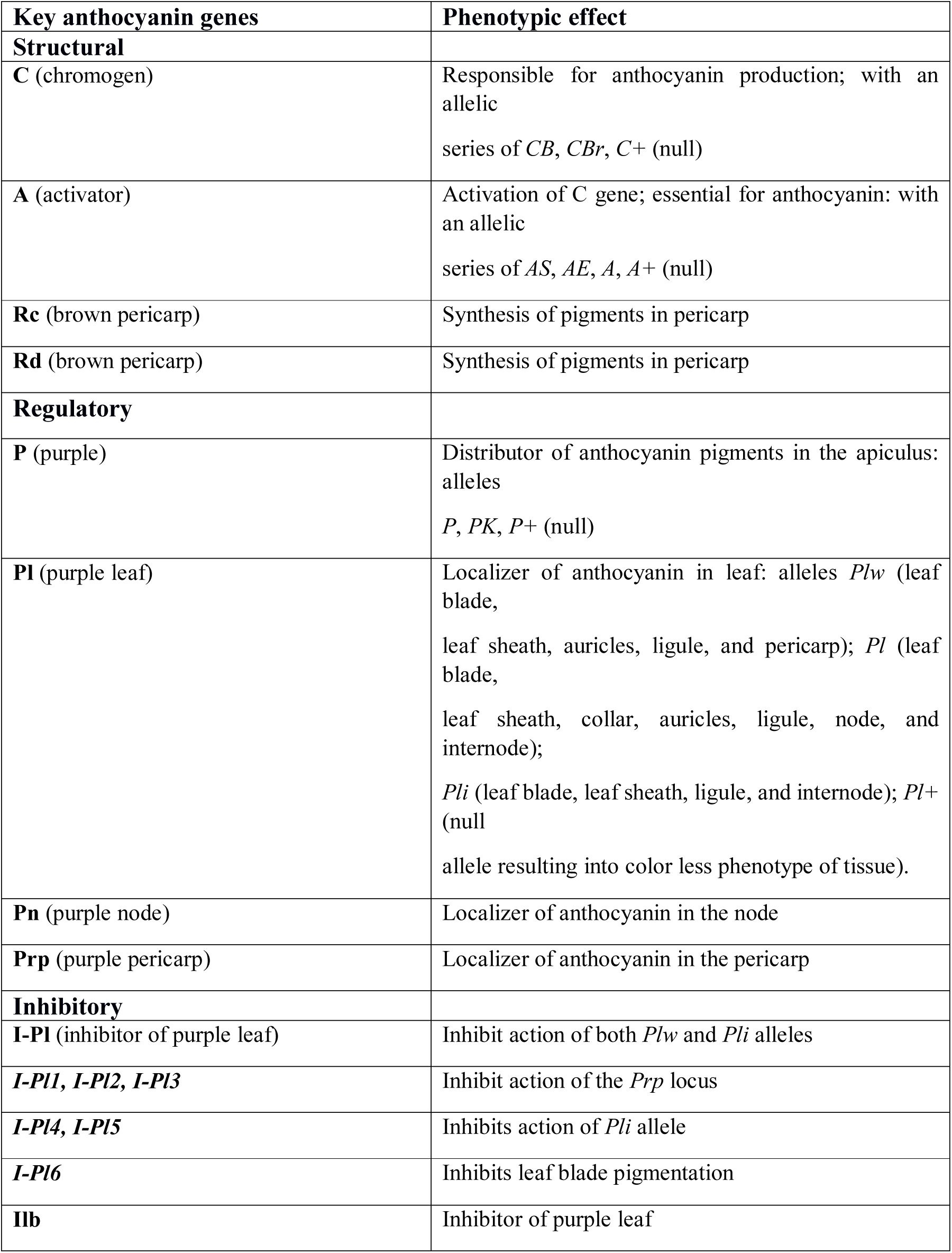
List of structural, regulatory and inhibitory genes involved anthocyanin pathway with their phenotypic effects.

Recently, Sun *et al.*, (2018) proposed a *C-S-A* gene model for rice hull pigmentation: *C1* that encodes a MYB transcription factor and acts as a color-producing gene, and *S1* that encodes a bHLH protein that functions in a tissue-specific manner. *C1* interacts with *S1* and activates expression of *A1*, which encodes a dihydroflavonol reductase (Sun *et al.*, 2018). We uncovered unique variants of various TFs such as MYB, bHLH, FAR1, NAC, WRKY, ERF, bZIP, bHLH, AP2 and others (Supplementary Table S14) responsible for different biological, molecular and cellular processes in PP. The role of bHLH, MYB, MYBAS1 TFs and other genes such F3H, Ra & Rc in regulating anthocyanins and other associated pathways responsible for PP color and texture (Supplementary Table S15) (Pireyre *et al.*, 2015) has been described here. In addition, as many as 96 novel variations in PP rice with known effects on morphological, physiological effects and stress response phenotypes are also reported.

Overall present study deals with whole genome variations in PP rice examined by identifying SNP and INDEL polymorphisms using 108 rice genome panel and SNP-Seek dataset. Base substitutions and distribution of DNA polymorphism over PP genome provides important insights into the molecular basis underlying phenotypic traits exhibited by the genotype. Further, the unique allele variations in different genes participating in flavonoid biosynthesis and anthocyanin pathways responsible for purple color in PP genome were revealed by comparative analysis. Unique SNPs identified in anthocyanin pathway occurring in both structural genes and regulatory transcription factors, will help to breed rice with high nutraceutical content, particularly, flavonoids that have antioxidant activity.

## Experimental Procedures

### Plant sample preparation and sequencing

Seeds of PP was germinated, and 15-day old seedlings were used for genomic DNA extraction using Qiagen DNeasy kit (Qiagen). Qubit 2.0 fluorometer (Thermo Fisher) was used to quantify and Nanodrop 2000 (Thermo Fisher) for quality check of the isolated DNA. The DNA was fragmented to 300bp size using a Covaris M220 focused ultrasonicator. The fragmented DNA was purified, and sequencing libraries were prepared using Illumina TruSeq DNA sample prep kit (Illumina Inc., USA) as per manufacturer’s specifications. The quantity and size distribution of the libraries were carried out using a Bioanalyzer 2100 (Agilent Technologies). The quantified libraries were sequenced on Illumina HiSeq-2000 platform (Illumina Technologies) by paired-end sequencing to generate 90-base long reads with an insert size of 200-350bp (Supplementary Figure S2). Standard Illumina pipeline was used to filter the raw data generated. To remove low-quality reads and reads containing adaptor/primer contamination, FASTQ files were further subjected to stringent quality control using NGS QC Toolkit (v2.3) (Patel and Jain, 2012).

### Mapping of PP genome

BWA software (v0.7) was used for the mapping of high-quality filtered reads against Nipponbare rice genome (MSU7) download from Rice Genome Annotation Project Database (http://rice.plantbiology.msu.edu/) (Li and Durbin, 2009; Kawahara *et al.*, 2013). Further, only uniquely aligned reads (with mapping quality ≥30 and minimum read depth 10) are considered in this analysis. Base quality score re-calibration and INDEL realignment were performed using Genome Analysis Toolkit (GATK, v2.1.13) and genome-wide coverage was estimated by Samtools (v0.1.16) (McKenna *et al.*, 2010; Li *et al.*, 2009).

### *De novo* assembly of unmapped PP genome and functional annotation of genes

Unmapped reads of PP genome were assembled using MaSuRCA (v3.2.1) assembler with default options (Zimin *et al.*, 2013). The assembled genome was masked using RepeatMasker (v4.0.6) with default parameters, using rice as a model (Smit *et al.*, 2015). Subsequently the masked genome was used in Augustus v3.2.1 using rice as model organism for gene prediction (Stanke *et al.*,2008). The annotations of identified genes were done using the Diamond program against NCBI NR, UniProt and Phytozome (v11.0) databases (Buchfink *et al.*, 2015).

### Identification and analysis of variants in PP genome

Minimum variant frequency of ≥90%, average base quality of the SNP ≥ 30 and minimum read depth of 10 were the stringent criteria followed to filter the identified SNPs and INDELs. If three or more SNPs were present in any 10-bp window, the SNPs and INDELs were filtered. Genome wide distribution of DNA polymorphisms was analysed by calculating their frequency at every100 kb interval on each rice chromosome. Circos was used to visualize the distribution of the SNPs and INDELs on rice chromosomes (Krzywinski *et al.*, 2009). Such distribution is assessed by integrating the position of DNA polymorphisms with GFF file containing rice genomic annotation. Customized Perl scripts were used to perform the genomic distribution and annotation of SNPs and INDELs. SnpEff (v3.1) tool was used to identify synonymous and nonsynonymous SNPs, and large-effect SNPs and INDELs (Cingolani *et al.*, 2012). Cut-off for number of nonsynonymous SNPs per Kb gene length was determined using Box and Whisker plot to identify the outlier genes.

### Comparative analysis of PP variations with diverse global rice lines collection

Initially, SNP dataset derived from alignment to IRGSP v1.0 was downloaded from SNP-seek database (http://snp-seek.irri.org/). Corresponding SNPs of PP were filtered as per SNP-seek database norms and compared using Bcftools to determine unique alleles. Identification of different transcripts associated with these variants were also performed. Variations in transcription factor genes were detected using PlantTFDB (Jin *et al.*, 2016). Moreover, comparative analysis of PP genome variations with 3024 rice genome collections available at Rice SNP-Seek Database was performed (Mansueto *et al.*, 2016).

### Annotation of variant-affected pathways and traits

All pathways associated with different variants have been annotated using rice metabolic pathway database genes (RiceCyc v3.3) using default parameters (Dharmawardhana *et al.*, 2013). Subsequently Q-TARO database (http://qtaro.abr.affrc.go.jp/) was used to identify variant-associated changes in morphological, physiological and resistance/tolerance traits (Yonemaru *et al.*, 2010; Yamamoto *et al.*,2012).

## Accession Number

Whole genome data of PP rice submitted in SRA database with bio project ID: PRJNA309223.

## Contributions

GT, SS, VBRL conceived the work and designed the experiments. RG, VBRL, BK, SPL, KAVSK, NC, LM performed *in silico* and *in vitro* experiments. RG, VBRL, GT, ARR analyzed the results. All contributed to writing the manuscript and discussed the results and commented on the manuscript.

## Compliance with Ethical Standards

This research does not perform any experiment on humans or animals. Data generated in this work was related to rice plant and submitted in SRA database (Acc No. PRJNA309223). Hence all the authors declare that there is no non-compliance with ethical standards.

## Supplementary Tables Information

**Supplementary Table S1.** List of annotated 1,239 gene sequences identified in PP unmapped genome.

**Supplementary Table S2.** Statistics of variant density across all chromosomes.

**Supplementary Table S3.** Repeats classes present in different variants falling into repeat regions of all chromosomes.

**Supplementary Table S4.** Different class of genes present in variant desert region on chromosome 5.

**Supplementary Table S5.** Gene summary for the variant desert region of chromosome 5.

**Supplementary Table S6.** Variant counts for the upstream (−1500 bp) region of the genes.

**Supplementary Table S7.** List of selected rice genomes for comparative analysis to identify unique variations in PP genome.

**Supplementary Table S8.** List of identified unique variants in PP genome after performing comparative analysis with 108 rice genomes.

**Supplementary Table S9.** List of identified unique variants in desert region of chromosome 5 after performing comparative analysis with 108 rice genomes.

**Supplementary Table S10.** Novel identified genes specific to PP genome and their associated traits.

**Supplementary Table S11.** List of 1,387 unique SNPs identified in PP genome after comparison with global collection of 3,023 rice lines.

**Supplementary Table S12.** Various RiceCyc pathways and associated transcripts and their different variants.

**Supplementary Table S13.** Summary of different variants involved in morphological, physiological and Resistance/tolerance traits.

**Supplementary Table S14.** Summary of the transcription factor with variations.

**Supplementary Table S15.** List of chromosome-wise identified variants and their associated genes information involved in anthocyanin pathway regulation.

## Supplementary Figures Information

**Supplementary Figure S1.**
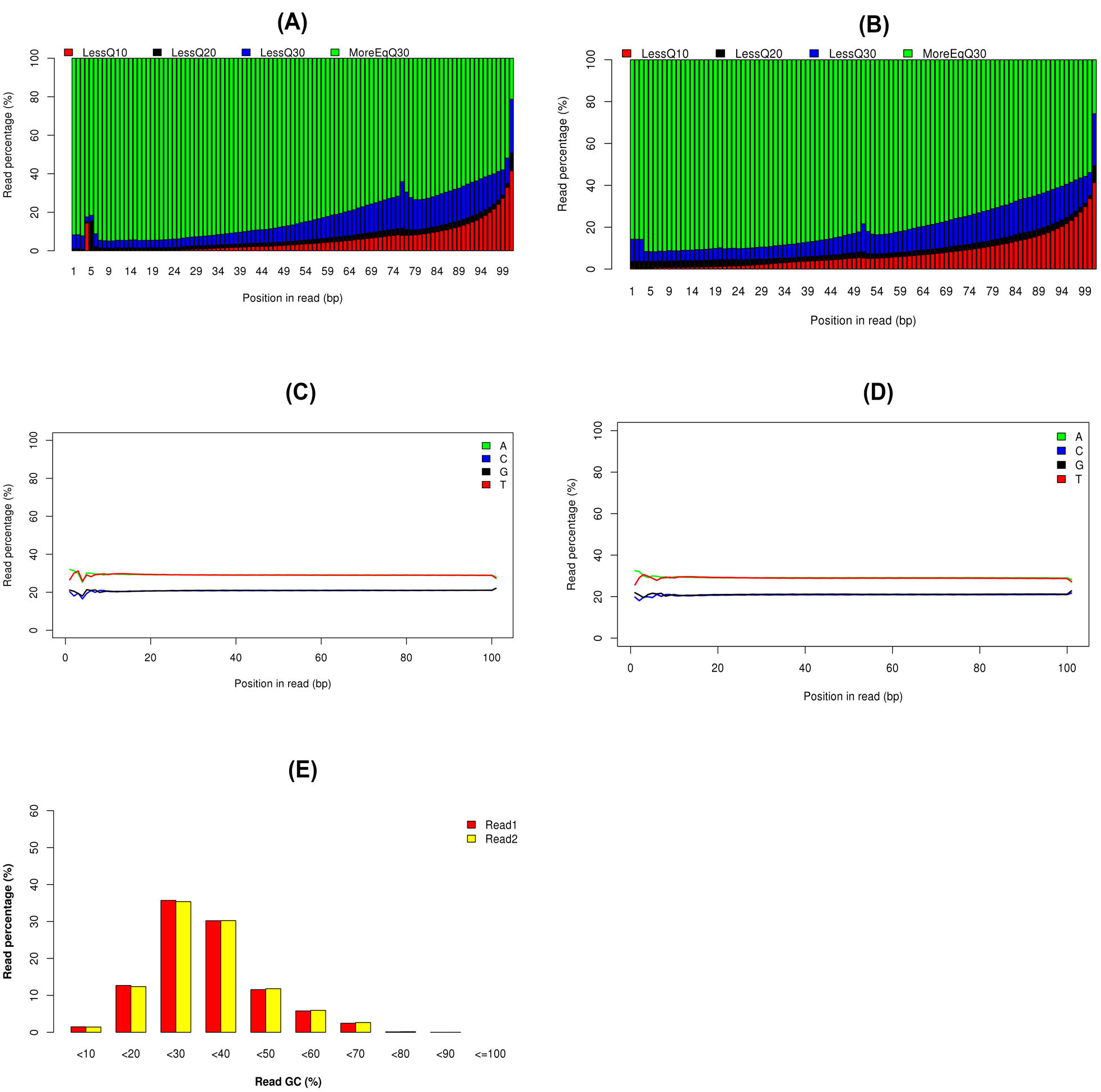
Sequencing quality scores of (A) read 1 (Forward) (B) read 2 (Reveres) reads, (C) & (D) indicates position-based quality of each nucleotide present in read 1 and read 2 respectively. (E) Shows the overall percentage of GC content *vs.* read percentages of both reads.

**Supplementary Figure S2.**
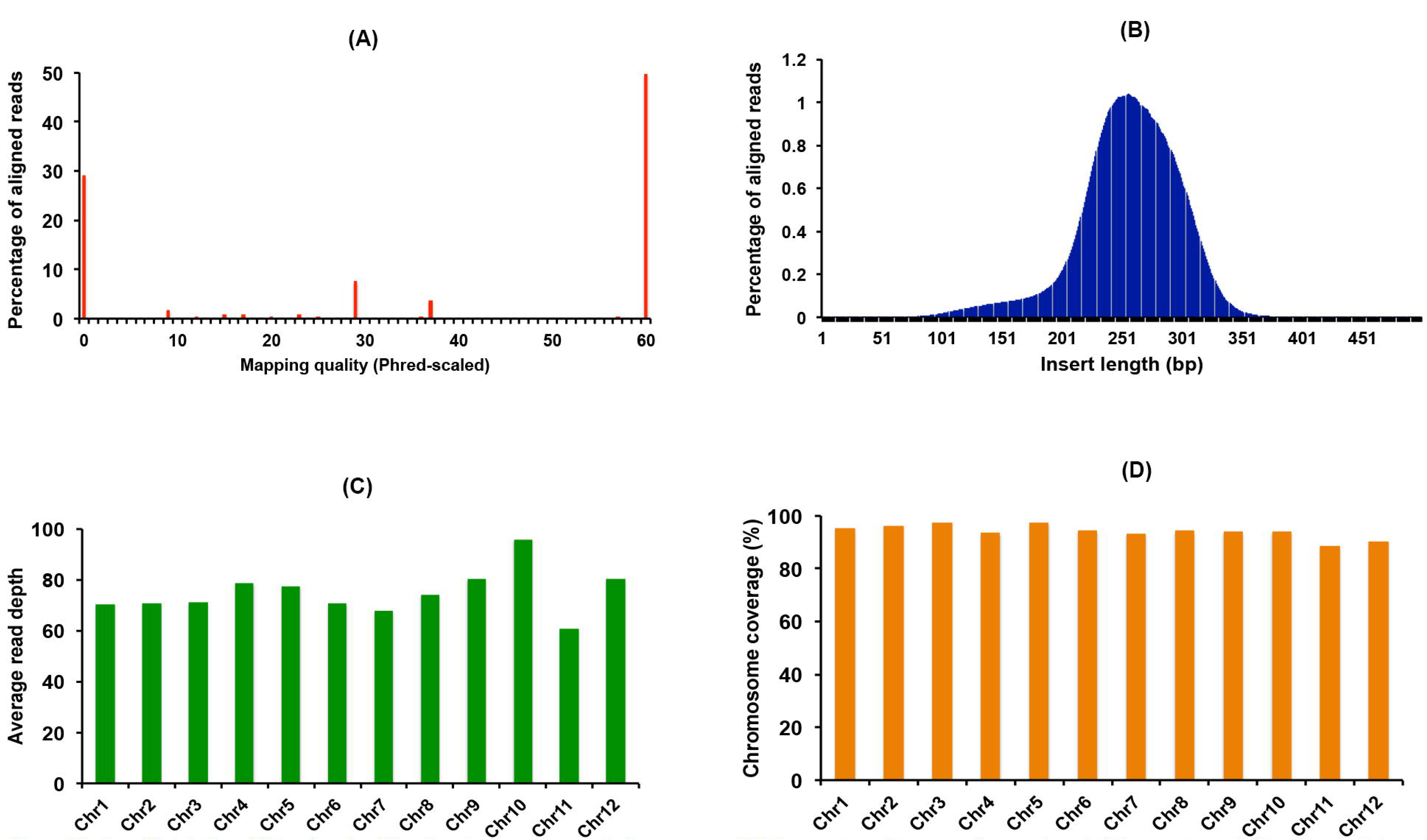
Overall statistics of filtered reads of Purpleputtu present over all chromosomes. (A) Represent quality score of mapped reads (B) percentage of aligned reads *Vs* insert length (C) Chromosome-wise average read depth and (D) Chromosome-wise coverage percentage.

**Supplementary Figure S3.**
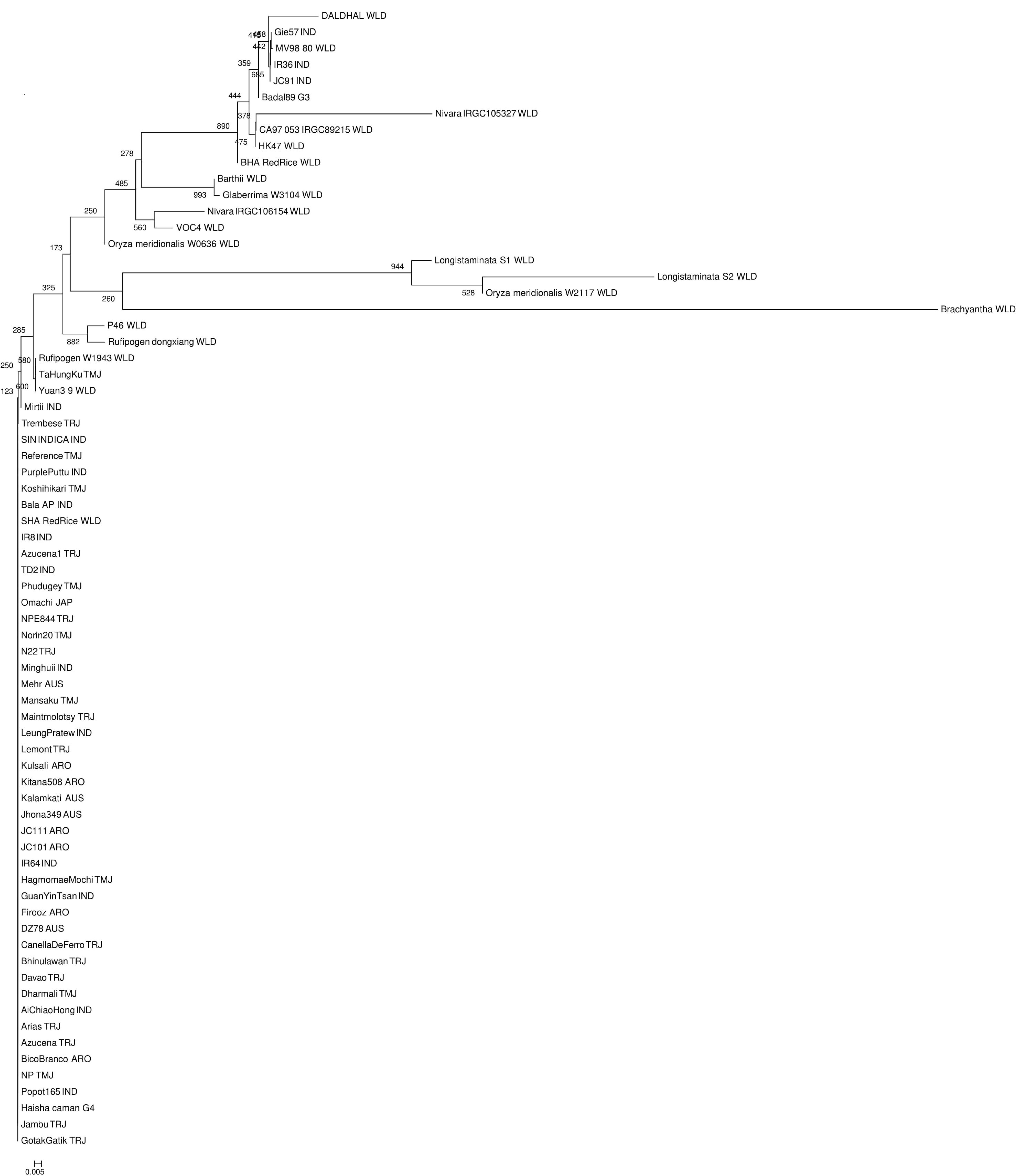
Phylogenetic tree diagram for Bh4 gene of PP rice with others rice cultivar Bh4 genes.

**Supplementary Figure S4.**
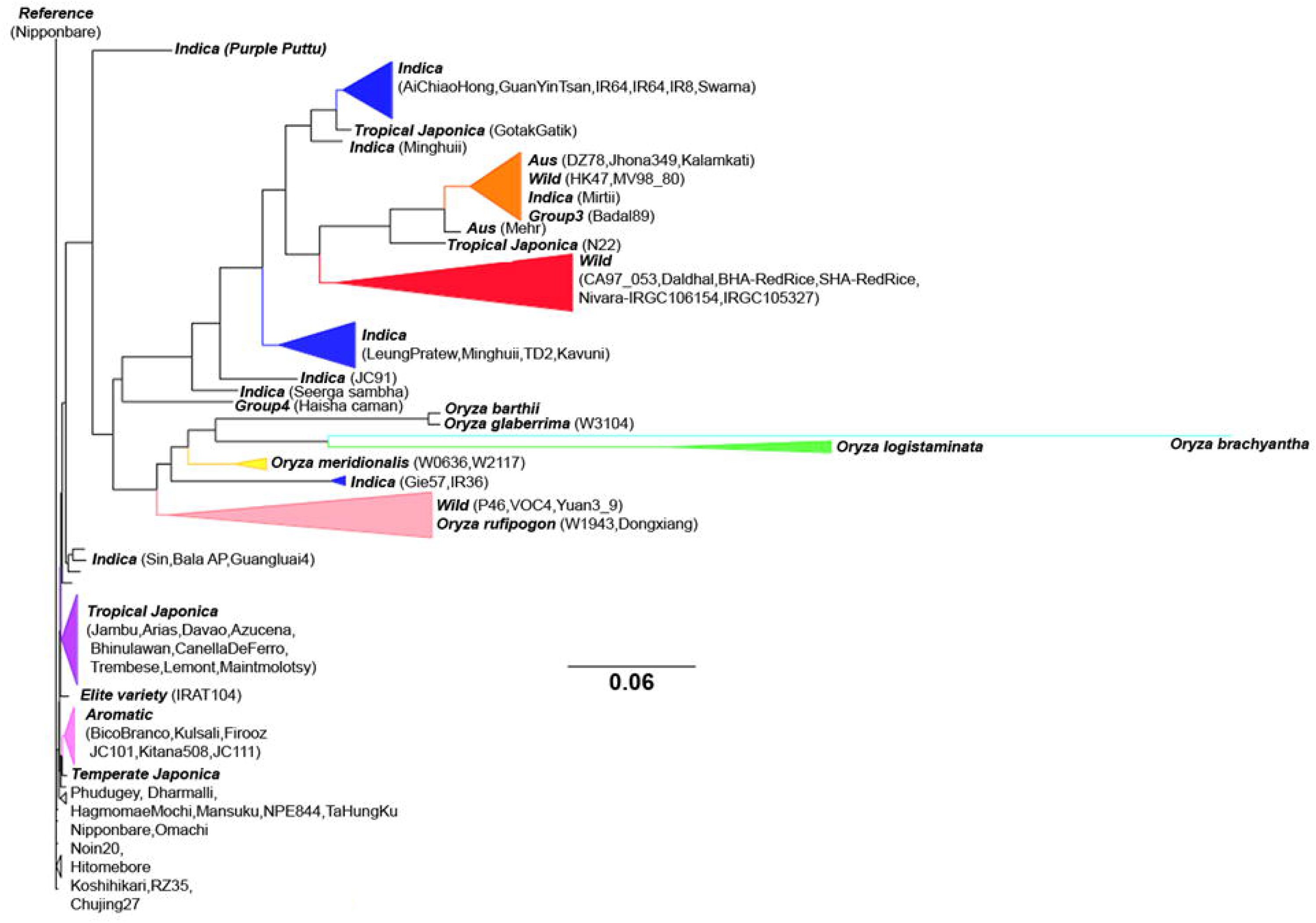
Dendrogram showing the Rc locus evolutionary divergence with other rice lines.

**Supplementary Figure S5.**
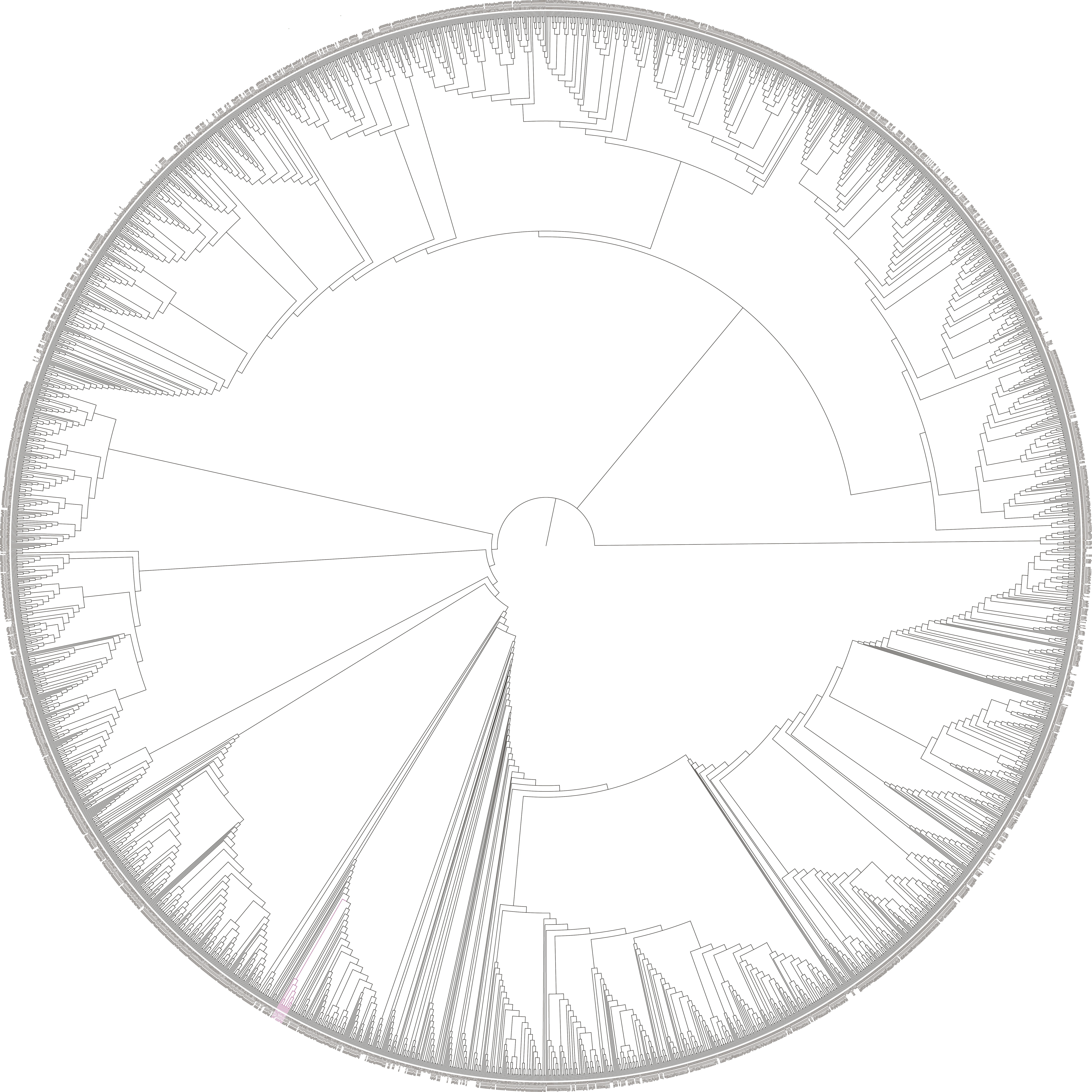
Dendrogram showing the divergence of PP with global collection of 3,023 rice lines.

